# Morphine differentially alters the synaptic and intrinsic properties of D1R- and D2R-expressing medium spiny neurons in the nucleus accumbens

**DOI:** 10.1101/858100

**Authors:** Dillon S. McDevitt, Benjamin Jonik, Nicholas M. Graziane

## Abstract

Exposure to opioids reshapes future reward and motivated behaviors partially by altering the functional output of medium spiny neurons (MSNs) in the nucleus accumbens shell. Here, we investigated how morphine, a highly addictive opioid, alters synaptic transmission and intrinsic excitability on dopamine D1-receptor (D1R) expressing and dopamine D2-receptor (D2R) expressing MSNs, the two main output neurons in the nucleus accumbens shell. Using whole-cell electrophysiology recordings, we show, that 24 h abstinence following repeated non-contingent administration of morphine (10 mg/kg, i.p.) in mice reduces miniature excitatory postsynaptic current (mEPSC) frequency and miniature inhibitory postsynaptic current (mIPSC) frequency on D2R-MSNs, with concomitant increases in D2R-MSN intrinsic membrane excitability. We did not observe any changes on synaptic or intrinsic changes on D1R-MSNs. Lastly, in an attempt to determine the integrated effect of the synaptic and intrinsic alterations on the overall functional output of D2R-MSNs, we measured the input-output efficacy by measuring synaptically-driven action potential firing. We found that both D1R-MSN and D2R-MSN output was unchanged following morphine treatment.

## Introduction

Exposure to opioids reshapes future reward and motivated behaviors partially by altering the functional output of medium spiny neurons (MSNs) in the nucleus accumbens shell, a brain region central to reward and motivation (Wolf, 2010;Graziane et al., 2016;Hearing et al., 2016;Scofield et al., 2016). MSNs receive glutamatergic excitatory input from the infralimbic prefrontal cortex, amygdala, hippocampus, and midline nuclei of the thalamus, while also receiving inhibitory input locally from interneurons or collateral projections from MSNs or from other brain regions including the ventral pallidum, lateral septum, periaqueductal gray, parabrachial nucleus, pedunculopontine tegmentum and ventral tegmental area (Sesack and Grace, 2010;Lalchandani et al., 2013;Salgado and Kaplitt, 2015;Dobbs et al., 2016;McDevitt and Graziane, 2018). The integration of these synaptic inputs along with the intrinsic excitably of medium spiny neurons (MSNs) are, in part, critically important for information transfer through the reward neurocircuit (Russo et al., 2010;Kourrich et al., 2015).

There are two main classes of MSNs in the accumbens shell; dopamine D1-receptor containing and dopamine D2-receptor containing MSNs (D1R-MSN and D2R-MSN, respectively). These cell-types not only differ in the dopamine receptor expressed, but also in their projection sites, peptidergic expression, and modulation of motivated behaviors (Hikida et al., 2010;Lobo et al., 2010;Smith et al., 2013;Koo et al., 2014;Al-Hasani et al., 2015;Creed et al., 2016;Heinsbroek et al., 2017;Tejeda et al., 2017;Castro and Bruchas, 2019). Recently, reports have demonstrated that exposure to morphine differentially alters excitatory glutamatergic transmission on both D1R- and D2R-MSNs in the accumbens shell (Graziane et al., 2016;Hearing et al., 2016;Hearing et al., 2018;Madayag et al., 2019). However, little is known regarding how exposure to morphine alters MSN cell-type specific inhibitory transmission and intrinsic membrane excitability, or how these synaptic and intrinsic factors integrate to drive future D1R-or D2R-MSN functional output. In an attempt to identify the effect of morphine exposure on D1R- and D2R-MSN functional output in the accumbens shell, we investigated how repeated exposure to morphine affected the integration of excitatory and inhibitory transmission, along with the intrinsic factors that drive membrane excitability. Finally, we assessed the integrated effect that synaptic and intrinsic factors had on the overall functional output of MSNs in the accumbens 24 h following morphine administration.

## 2. Materials and methods

### 2.1. Animals

All experiments were done in accordance with procedures approved by the Pennsylvania State University College of Medicine Institutional Animal Care and Use Committee. Cell-type specific D1R- or D2R-MSN recordings were made using male and female B6 *Cg-Tg* (*Drd1a*-tdTomato) line 6 Calak/J hemizygous mice, a bacterial artificial chromosome (BAC) transgenic mouse line initially developed in the laboratory of Dr. Nicole Calakos at Duke University, aged 5-10 weeks (Ade et al., 2011) (JAX stock #16204). Given that in this transgenic mouse line, D1R-MSNs are fluorescently labeled, D2R-MSNs were identified based on the lack of fluorescence, cell size, and electrophysiological characteristics, including capacitance and membrane resistance (**Table I**), as previously published (Graziane et al., 2016). Additionally, as elegantly stated previously (Willett et al., 2019), unlabeled MSNs in the *Drd1a*-tdTomato line in adult mice nearly exclusively compromise *Drd2*-positive MSNs and to a lesser extent MSNs expressing both D1R and D2R (D1R/D2R-MSNs) (1.6%) (Ade et al., 2011;Enoksson et al., 2012;Thibault et al., 2013). Thus, we refer to all unlabeled MSNs from the *Drd1a*-tdTomato line as D2R-MSNs, but with the full acknowledgement that we are also likely sampling from D1R/D2R-MSNs, but to a much lesser degree (Bertran-Gonzalez et al., 2008;Ade et al., 2011). Mice were singly-housed and maintained on a regular 12 hour light/dark cycle (lights on 07:00, lights off 19:00) with *ad libitum* food and water.

**Table I.** Cell Properties from electrophysiological assessments.

|  | **Saline** |  | **Morphine** |  |
| --- | --- | --- | --- | --- |
|  | D1R-MSN | D2R-MSN | D1R-MSN | D2R-MSN |
| **Membrane Capacitance (pF)** | 81.11±2.11<br>(71) | 75.83±2.56<br>(64) | 82.01±2.48<br>(72) | 76.93±1.89<br>(78) |
| **Membrane Resistance (MΩ)** | 250.5±9.09<br>(71) | 256.1±12.34<br>(64) | 246.6±11.88<br>(72) | 280.3±10.8<br>(78) |
| **Tau (ms)** | 1.53±0.07<br>(21) | 1.42±0.09<br>(16) | 1.49±0.08<br>(21) | 1.49±0.12<br>(16) |

Number of cells (n). Tau values provided by Axon software. They were unavailable in Sutter Software, which results in a lower n.

### 2.2. Drugs

(−)-morphine sulfate pentahydrate was provided by the National Institute on Drug Abuse Drug Supply Program. NBQX and AP5 were purchased from Tocris Biosciences. Picrotoxin was purchased from Sigma Aldrich. Tetrodotoxin (TTX) was purchased from Enzo.

### 2.3. Repeated systemic injections of saline or morphine

Before drug administration, mice were allowed to acclimate to their home cages for >5d. For drug treatment, we used a 5d repeated drug administration procedure (Huang et al., 2009;Graziane et al., 2016). In all electrophysiological experiments, once per d for 5d, mice were taken out of the home cages for an intraperitoneal (i.p.) injection of either (−)-morphine sulfate pentahydrate (10mg/kg in 0.9% saline) or the same volume of 0.9% saline, and then placed back to the home cage at ~Zeitgeber time (ZT) 2 (ZT0=lights on, ZT12=lights off). Animals were randomly selected for each drug treatment. Morphine- or saline-treated animals were then used for electrophysiological recordings ~24h following the last injection.

### 2.4. Acute Brain Slice Preparation

At ~ZT time 2, mice were deeply anesthetized with isoflurane and cardiac perfused with an ice-cold NMDG-based cutting solution containing (in mM): 135 N-methyl-d-glutamine, 1 KCl, 1.2 KH_2_PO_4_, 0.5 CaCl_2_, 1.5 MgCl_2_, 20 choline-HCO_3_, and 11 glucose, saturated with 95%O_2_/5%CO_2_, adjusted to a pH of 7.4 with HCl, osmolality adjusted to 305 mmol/kg. Following perfusion, mice were decapitated and brains were rapidly removed. 250 μm coronal brain slices containing the nucleus accumbens shell were prepared via a Leica VT1200s vibratome in 4°C NMDG cutting solution. Following cutting, slices were allowed to recover in artificial cerebrospinal fluid (aCSF) containing (in mM): 119 NaCl, 2.5 KCl, 2.5 CaCl_2_, 1.3 MgCl_2_, 1 NaH_2_PO_4_, 26.2 NaHCO_3_, and 11 glucose, osmolality of 290 mmol/kg, at 31°C for 30 minutes followed by 30 minutes at 20-22°C prior to recording. After a one hour recovery period, slices were kept at 20-22°C for the rest of the recording day.

### 2.5. Electrophysiology

Whole-cell recording. All recordings were made from the nucleus accumbens shell between Bregma 1.7 mm and 0.86 mm (Paxinos and Franklin, 2004). Slices were transferred to a recording chamber and neurons were visualized using infrared differential interference contrast microscopy. During recording, slices were superfused with aCSF at room temperature. For intrinsic membrane excitability experiments, recording electrodes (2-5 MΩ; borosilicate glass capillaries (WPI #1B150F-4) pulled on a horizontal puller from Sutter Instruments (model P-97)) were filled with a potassium-based internal solution containing (in mM): 130 KMeSO_3_, 10 KCl, 10 HEPES, 0.4 EGTA, 2 MgCl_2_-6H_2_0, 3 Mg-ATP, 0.5 Na-GTP, pH 7.2-7.4, osmolality=290 mmol/kg (Wescor Vapro Model 5600, ElitechGroup). Resting membrane potential was recorded immediately following break-in. Before beginning the protocol, cells were adjusted to a resting membrane voltage of −80mV. This typically was achieved with less than 30 pA current injection, and cells were discarded if the current needed to adjust the cell to −80 mV was greater than 50 pA. A current step protocol consisting of 600 ms steps ranging from −200 to +450 pA in 50 pA increments was carried out with a 20 s intra-sweep interval. The number of action potentials observed at each current step was recorded.

For synaptically-driven action potential experiments or rheobase/chronaxie measurements, a stimulation electrode (size, 2.5–3 MΩ), filled with aCSF, was placed 100 μm from the recorded neuron along the same z plane in three dimensional space. Recordings were performed using KMeSO_3_ as described above. The resting membrane potential was not adjusted, enabling neurons to fire action potentials. The average membrane potential during electrophysiology recordings was −85.3±0.78 mV, which deviated by 4.39±0.40 mV (n=50) throughout the entirety of the experiment. For synaptically-driven action potential experiments, a 10 Hz stimulus with a stimulus duration of 0.25 ms and stimulus strength ranging from 0-100 μAmps of current, with an interval of 5 μAmps, was applied through the stimulating electrode. For each current, this procedure was repeated three times and the average number of action potentials/10 Hz stimulus was recorded. Rheobase/chronaxie measurements were made by varying the stimulus duration from 2-0.2 μAmps and injecting current at each duration until an action potential was evoked from the recorded neuron. The stimulus duration was plotted over the current which elicited an action potential. The rheobase was calculated as the plateau of a two-phase decay nonlinear regression curve fit. The chronaxie was calculated, using GraphPad Prism software, as the duration corresponding to 2x the rheobase, by solving for x in the equation, rheobase*2=rheobase + SpanFast*exp(-KFast*x) + SpanSlow*exp(-KSlow*x).

For excitatory/inhibitory ratio (E/I) experiments (Liu et al., 2016), recording electrodes (2-5 MΩ) were filled with a cesium-based internal solution (in mM): 135 CsMeSO3, 5 CsCl, 5 TEA-Cl, 0.4 EGTA (Cs), 20 HEPES, 2.5 Mg-ATP, 0.25 Na-GTP, 1 QX-314 (Br), pH 7.2-7.4, osmolality=290 mmol/kg. This internal solution was selected i) to isolate synaptically-evoked currents (cesium and QX-314 block voltage-gated K^+^ and Na^+^ channels, respectively) and ii) to measure the E/I ratios at physiologically relevant ionic driving forces while MSNs were voltage clamped at −70 mV (−70 mV is similar to the membrane potential of MSNs during synaptically-driven action potential and rheobase/chronaxie measurements, which were performed in current clamp) (using this internal solution the reversal potential for γ-aminobutyric acid_A_ (GABA_A_) receptor/glycine receptors (receptors likely mediating inhibitory postsynaptic currents (IPSCs) and AMPA/kainate receptors (receptors mediating excitatory postsynaptic currents (EPSCs)) is ~-60 mV and ~0 mV, respectively). To evoke postsynaptic currents, presynaptic afferents were stimulated via a constant-current stimulator (Digitimer) using a monopolar stimulating electrode (glass pipette filled with aCSF) at 0.1 Hz with 0.1 ms stimulus duration. Cells were held at −70 mV for the entirety of the experiment. Once a stable baseline was observed near 200 pA of current, 50 traces were recorded. Following this, NBQX (2 μM) and AP5 (50 μM) were bath applied to isolate inhibitory ionotropic receptor-mediated currents. The drug was allowed to wash on, and 50 more sweeps were recorded. The AMPA/kainate receptor-mediated current was then obtained via digital subtraction of the inhibitory ionotropic receptor-mediated current from the mixed current. The E/I ratio was then calculated by taking the peak amplitude of the AMPA/kainate receptor-mediated current divided by the peak amplitude of the inhibitory ionotropic receptor-mediated current in male or female mice.

E-I balance assessments investigating temporal relationships between excitatory and inhibitory current were carried out in male mice by measuring spontaneous events using cesium based internal solution (see recipe above) and aCSF. Neurons were held at −30 mV in order to elicit inward excitatory current and outward inhibitory current, as done previously (Zhou et al., 2009). Recordings lasted three minutes and analysis was performed using MiniAnalysis software. A computer program built in Visual Studio was used to calculate the inter-event intervals of sEPSC and sIPSCs.

Miniature excitatory or inhibitory postsynaptic current (mEPSC or mIPSC, respectively) recordings were performed in the presence of tetrodotoxin (1 μM), a Na^+^ channel blocker. mEPSCs were recorded in the presence of picrotoxin (100 μM) and mIPSCs were recorded in the presence of NBQX (2 μM). For mEPSC recordings, recording electrodes (2-5 MΩ) were filled with cesium-based internal solution as described above. For mIPSC recordings, recording electrodes (2-5 MΩ) were filled with high chloride cesium-based internal solution (in mm): 15 CsMeSO3, 120 CsCl, 8 NaCl, 0.5 EGTA (Cs), 10 HEPES, 2.0 Mg-ATP, 0.3 Na-GTP, 5 QX-314 (Br), pH 7.2-7.4, osmolality=290 mmol/kg. High chloride cesium-based internal solution was used for mIPSC recordings so that mIPSCs could be detected in neurons voltage clamped at −70 mV (γ-aminobutyric acid_A_ (GABA_A_) receptor/glycine receptor reversal potential=~0 mV). Events during a stable 10 min period were analyzed using Sutter software (Pernia-Andrade et al., 2012). Decay tau corresponds to the time constant of decay time, which equals the 10-90% decay time. The rise time equals 10-90% rise time.

All recordings were performed using either an Axon Multiclamp 700B amplifier or Sutter Double IPA, filtered at 2-3 kHz, and digitized at 20 kHz. Series resistance was typically 10-25 MΩ, left uncompensated, and monitored throughout. For all voltage clamp recordings, cells with a series resistance variation greater than 20% were discarded from analysis. For all current clamp recordings, cells with a bridge balance that varied greater than 20% during the start and end of recordings were discarded from analysis.

### 2.6. Statistical Analysis

All results are shown as mean±SEM. Each experiment was replicated in at least 3 animals. No data points were excluded. Sample size was presented as n/m, where “n” refers to the number of cells and “m” refers to the number of animals. Statistical significance was assessed in GraphPad Prism software using a one- or two-way ANOVA with Bonferroni’s correction for multiple comparisons in order to identify differences as specified. F values for two-way ANOVA statistical comparisons represent interactions between variables unless otherwise stated. Two-tail tests were performed for all studies. Our goal, a priori, was to examine pairwise comparisons between drug treatment and cell type combinations regardless if the interaction effect between drug treatment and cell type was strong. Thus, prior to analysis, we created all possible independent groups based on drug treatment and cell type combinations and performed a one-way ANOVA with pairwise comparisons. The results from these pairwise comparisons from this one-way ANOVA would be equivalent to performing a two-way ANOVA with an interaction term (drug treatment, cell type, drug treatment*cell type interaction) and then performing post-hoc pairwise comparisons on the interaction term from the two-way ANOVA model.

## 3. Results

### 3.1. Morphine reduces synaptic transmission on D2R-MSNs

Previously, it was found that exposing mice to a dosing regimen (i.p. 10 mg/kg per d for 5 d, 1-d forced abstinence) that induces locomotor sensitization and conditioned place preference generates silent synapse expression preferentially on D2R-MSNs, but not D1R-MSNs, via removal of AMPA receptors from mature synapses (Graziane et al., 2016). The removal of AMPA receptors from the synapse is expected to change the number of release sites (n) when AMPA receptor-mediated transmission is the readout (Hanse et al., 2013), which results in a change in frequency of quantal events (Kerchner and Nicoll, 2008). Based on this, we assessed morphine-induced quantal changes in D1R- or D2R-MSN synaptic transmission by measuring mEPSCs. We found that 24 h following repeated morphine treatment, D1R-MSNs showed no changes in mEPSC amplitude (Bonferroni post-test, p>0.999) (Figs. 1A, B, D), and this was also observed on D2R-MSNs (Bonferroni post-test, p>0.999) (Figs. 1A, C, D). Furthermore, analysis of mEPSC frequency following morphine exposure showed no effect on D1R-MSNs (Bonferroni post-test, p=0.19) (Figs. 1E and G). However, post-hoc analysis revealed a significant difference between mEPSC frequency following morphine exposure on D2R-MSNs (Bonferroni post-test, p=0.01) (Figs. 1F and G). Lastly, we analyzed the receptor rise time and decay tau of mEPSCs in order to measure whether the significant effects observed were potentially mediated by changes in AMPA/kainate receptor kinetics. We found, in all groups, the receptor kinetics, rise time and decay tau, remained unchanged (Rise time: F_(3,42)_=0.371, p=0.77; One-way ANOVA; decay tau: F_(3,42)_=0.290, p=0.83; One-way ANOVA) (Fig. 1H and I). Based on previously published findings (Graziane et al., 2016), it is likely that the observed morphine-induced decreases in D2R-MSN mEPSC frequency are caused by a reduction in the number of release sites (n) due to morphine-induced AMPA receptor removal from mature D2R-MSN synapses.

**Figure 1.**
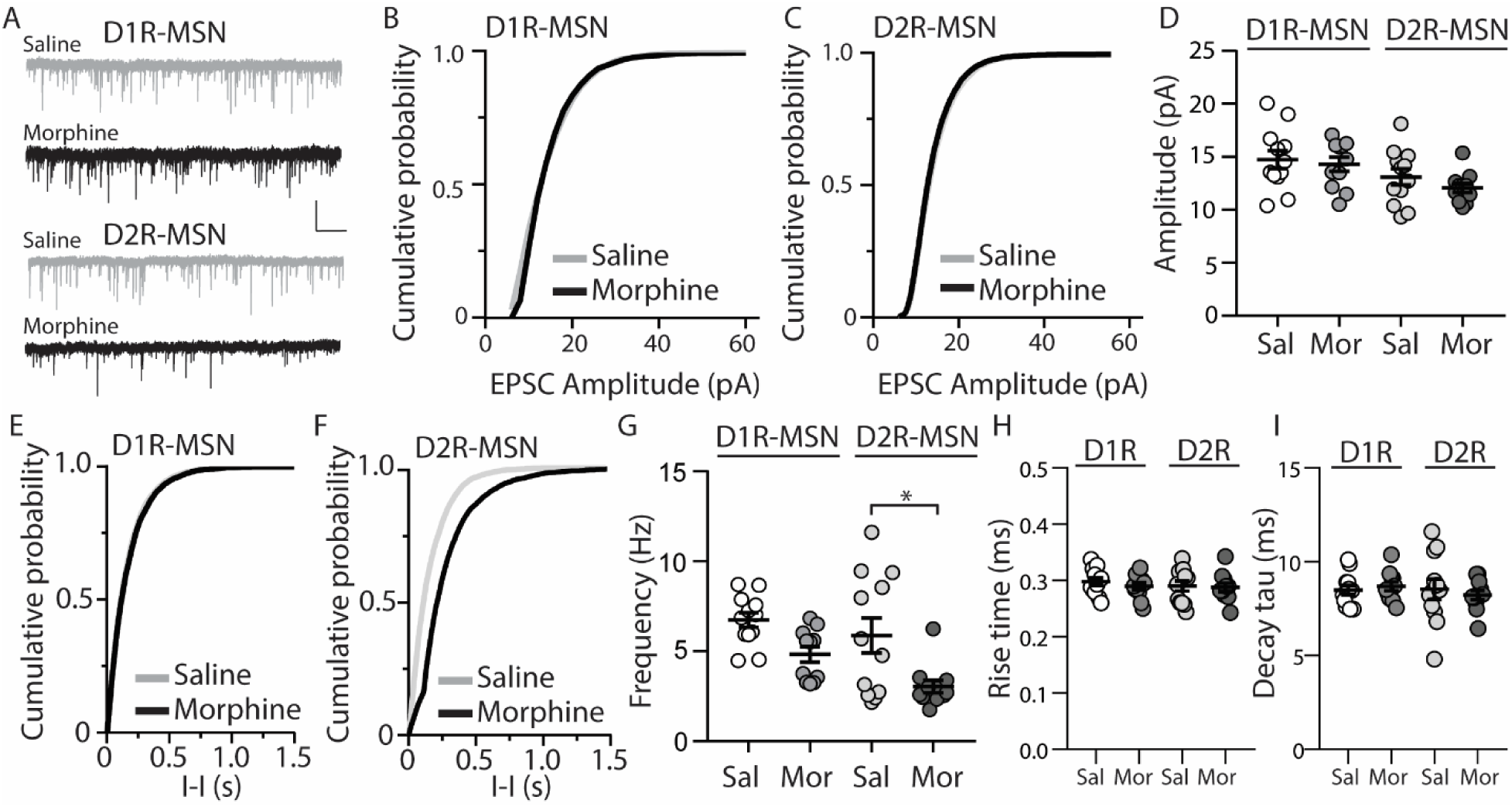
Repeated morphine administration reduces mEPSC frequency on D2R-MSNs on abstinence day 1. (A) Representative traces showing mEPSCs recorded from D1R- or D2R-MSNs from animals treated with saline or morphine (10 mg/kg, i.p.). Scale bar: 20 pA, 1s. (B and C) Cumulative plot of a representative neuron showing the distribution of mEPSC amplitudes recorded from D1R-MSNs (B) or D2R-MSNs (C) in animals treated with saline or morphine. (D) Summary graph showing the average mEPSC amplitude recorded D1R- or D2R-MSNs following saline or morphine treatment (F_(3,42)_=2.92, p=0.045; One-way ANOVA. Bonferroni post-test, D1R-MSN: saline versus morphine, p>0.999; D2R-MSN: saline versus morphine, p>0.999). (E and F) Cumulative plot of a representative neuron showing the distribution of mEPSC inter-event intervals (I-I) recorded from D1R-MSNs (E) or D2R-MSNs (F) in animals treated with saline or morphine. (G) Summary graph showing the average mEPSC frequency recorded from D1R- or D2R-MSNs following saline or morphine treatment (F_(3,42)_=6.73, p=0.0008; One-way ANOVA with Bonferroni post-test; *p<0.05, **p<0.01). (H) Summary graph showing the average rise time of mEPSC recorded from D1R- or D2R-MSNs following saline or morphine treatment. (I) Summary graph showing the average decay tau of mEPSC recorded from D1R- or D2R-MSNs following saline or morphine treatment. Circle=neuron.

The functional output of MSNs in the nucleus accumbens relies upon the integration of excitatory and inhibitory synaptic transmission (Plenz and Kitai, 1998;Wickens and Wilson, 1998;Wolf et al., 2005;Otaka et al., 2013). To measure whether morphine-induced changes in mEPSC frequency on D2R-MSNs are sufficient to impact the excitatory and inhibitory balance of synaptic input, we measured the ratio of excitatory ionotropic receptor-mediated current to inhibitory ionotropic receptor-mediated current (E/I ratio) following an electrically evoked stimulus while MSNs were voltage-clamped at −70 mV. We found that 24 h post morphine treatment the E/I ratios were unchanged on D1R- or D2R-MSNs (F_(3,37)_=1.27, p=0.30; One-way ANOVA) (Fig. 2A and B). Given that the temporal integration of excitatory and inhibitory synaptic transmission regulates neuronal activity (Wehr and Zador, 2003;Higley and Contreras, 2006;Okun and Lampl, 2008;Hiratani and Fukai, 2017;Roland et al., 2017;Bhatia et al., 2019), we investigated whether morphine administration altered the temporal relationship between excitation and inhibition on D1R- or D2R-MSNs. In order to test this, we recorded spontaneous EPSCs (sEPSCs) and sIPSCs while voltage clamping D1R- or D2R-MSNs at −30 mV, which enabled us to simultaneously detect EPSCs (inward current with a reversal at ~0 mV (Lee et al., 2013)) and IPSCs (outward current with a reversal potential at ~-60 mV; **Table II**) within each neuron, as previously demonstrated (Zhou et al., 2009). Measuring spontaneous activity was chosen in order to sample both action potential mediated and non-action potential mediated events, encompassing synaptic populations sampled during evoked stimulation or miniature postsynaptic current recordings, respectively (He et al., 2018). With this approach, we were able to measure the temporal relationship between sEPSCs and sIPSCs as well as the balance of excitatory to inhibitory transmission on D1R- or D2R-MSNs (Fig. 2C). Our results show that morphine exposure did not alter the temporal relationship between excitatory and inhibitory events as we did not observe any changes in the inter-event interval between sEPSCs to sIPSCs (F_(3,27)_=0.198, p=0.90, one-way ANOVA) (Fig. 2D) or from sIPSCs to sEPSCs (F_(3,27)_=0.072, p=0.97, one-way ANOVA) (Fig. 2E). Additionally, we did not observe any morphine-induced changes in the excitation to inhibition balance measured by taking the sEPSC/sIPSC frequency ratio on D1R- or D2R-MSNs (F_(3,27)_=0.339, p=0.80, one-way ANOVA) (Fig. 2F), suggesting that the relationship between spontaneous postsynaptic excitatory and inhibitory currents within a neuron are unaffected by morphine treatment, despite the observed changes in mEPSC frequency.

**Figure 2.**
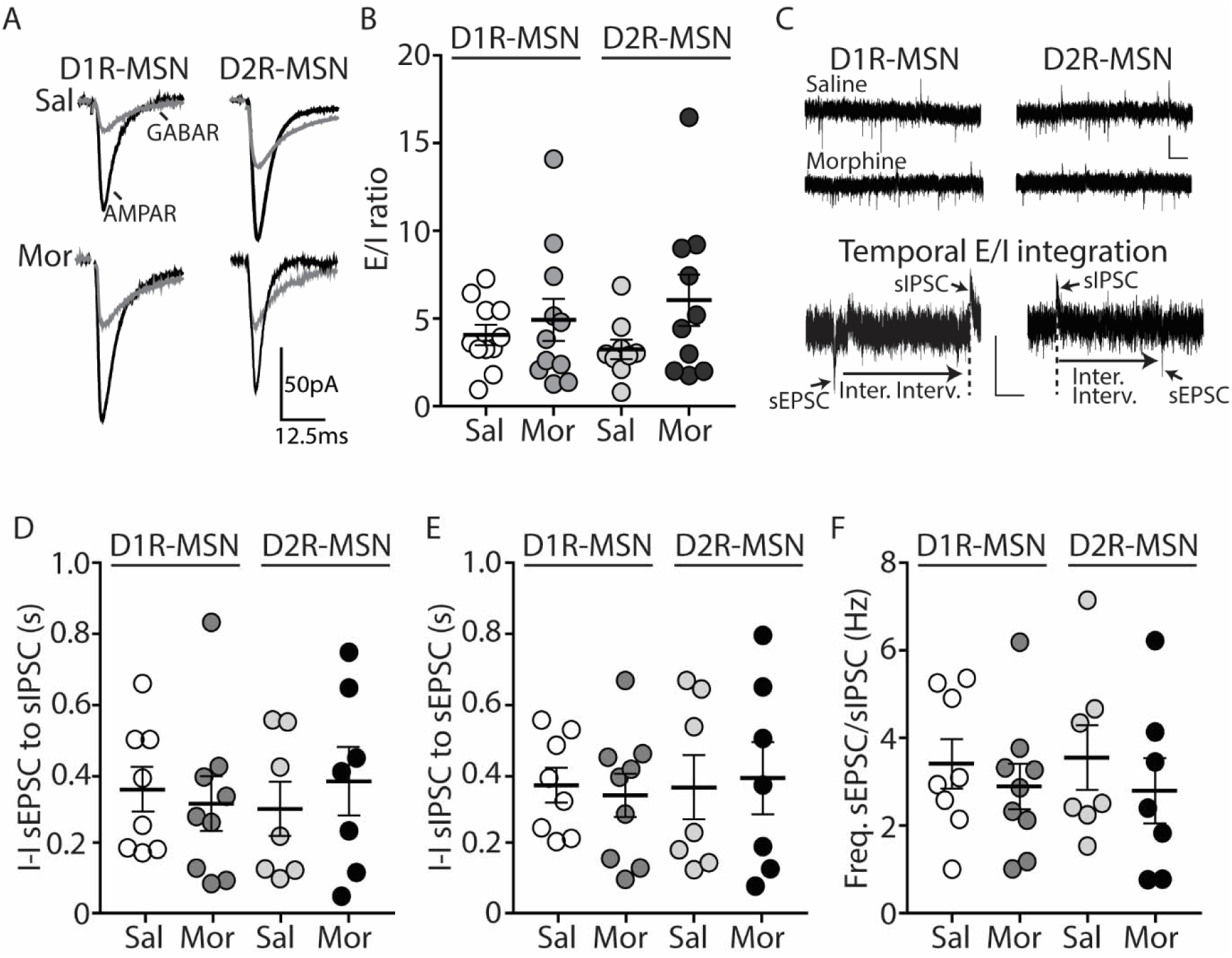
Short-term abstinence from *in vivo* morphine treatment has no effect on the evoked excitatory/inhibitory (E/I) ratio and does not alter the temporal relationship between spontaneous EPSCs (sEPSCs) and IPSCs (sIPSCs) on D1R- or D2R-MSNs in the nucleus accumbens shell. (**A**) Representative traces showing evoked AMPA receptor (AMPAR)- and GABA receptor (GABAR)-mediated currents on D1R- or D2R-MSNs 24 h following repeated saline or morphine treatments. Neurons were held at −70 mV. (**B**) Summary graph showing the E/I ratio of evoked currents on D1R- or D2R-MSNs 24 h following repeated saline (sal) or morphine (mor) (10 mg/kg, i.p.) treatments. There were no significant differences between groups in male or female mice. (**C**) Representative traces showing spontaneous EPSCs (inward current) and IPSCs (outward current) when D1R- or D2R MSNs were held at −30 mV 24 h following *in vivo* morphine treatment. Scale bars: 20 pA, 0.5 s (Lower). Electrophysiological recordings in whole-cell patch clamp configuration showing the inter-event intervals (inter. interv.) of sEPSCs to sIPSCs (left) or sIPSCs to sEPSCs (right) in a MSN held at −30 mV. Scale bars: 20 pA, 0.125 s (**D**) Summary graph showing no significant changes in the inter-event interval (I-I) between sEPSCs and sIPSCs on D1R- or D2R-MSNs 24 following repeated saline or morphine administration in male mice. (**E**) Summary graph showing no significant changes in the interevent interval (I-I) between sIPSCs and sEPSCs on D1R- or D2R-MSNs 24 following repeated saline or morphine administration in male mice. (**F**) Summary graph showing that morphine exposure had no effect on the frequency ratio of sEPSC to sIPSC events within D1R- or D2R-MSNs in male mice.

**Table II.** Calculated Cl- reversal potential for D1R- or D2R-MSNs in the nucleus accumbens shell. Cl-reversal potential was calculated in the whole-cell patch-clamp configuration using cesium methanesulfonate internal solution with bath application of aCSF+NBQX (2μM) and AP5 (50 μM). Whole-cell patch clamp configuration was used to mimic the approach used in spontaneous EPSC and IPSC recordings (Fig. 2). The values were corrected with a junction potential of 10.4 mV (V_m_=V_p_-V_L_ where V_m_=the membrane voltage, V_p_=the calculated voltage, and V_L_ is the voltage of the liquid junction potential) (Figl et al., 2003). N/m=number of cells/number of animals.

| Saline |  | Morphine |  |
| --- | --- | --- | --- |
| D1R-MSN | D2R-MSN | D1R-MSN | D2R-MSN |
| -59.09±1.2 mV;<br>n=10/7 | -60.98±1.5 mV; n=8/5 | -59.19±1.7 mV; n=5/4 | -60.94±1.0 mV; n=4/4 |

Because E/I ratios are dependent upon changes in excitatory and/or inhibitory synaptic transmission, we next investigated whether inhibitory transmission on D1R- or D2R-MSNs was altered 24 h following morphine treatment, by measuring miniature inhibitory postsynaptic currents (mIPSCs) (Fig. 3A). First, we measured mIPSC amplitude on D1R- or D2R-MSNs. We found that following morphine treatment, there was no significant change in the mIPSC amplitude on D1R-MSNs (Bonferroni post-test, p>0.999) (Figs. 3B and D) or on D2R-MSNs (Bonferroni post-test, p>0.999) (Figs. 3C and D). Furthermore, when measuring the mIPSC frequency, our data revealed no significant morphine-induced change on D1R-MSNs (Bonferroni post-test, p=0.949) Figs. 3E-G). However, morphine abstinence elicited a significant decrease in mIPSC frequency on D2R-MSNs (Bonferroni post-test, p<0.0001) (Figs. 3F and G). We also found that basal levels of mIPSC frequency were significantly greater on D2R-MSNs compared to D1R-MSNs (Bonferroni post-test, p=0.02). Lastly, to measure whether inhibitory ionotropic receptor kinetics were potentially a factor in the observed changes, we measured mIPSC rise time and decay tau (Fig. 3H and I). We found, in all groups, the receptor kinetics, rise time and decay tau, remained unchanged (rise time: F_(3,65)_=1.69, p=0.18; decay tau: F_(3,65)_=1.62, p=0.19; one-way ANOVA).

**Figure 3.**
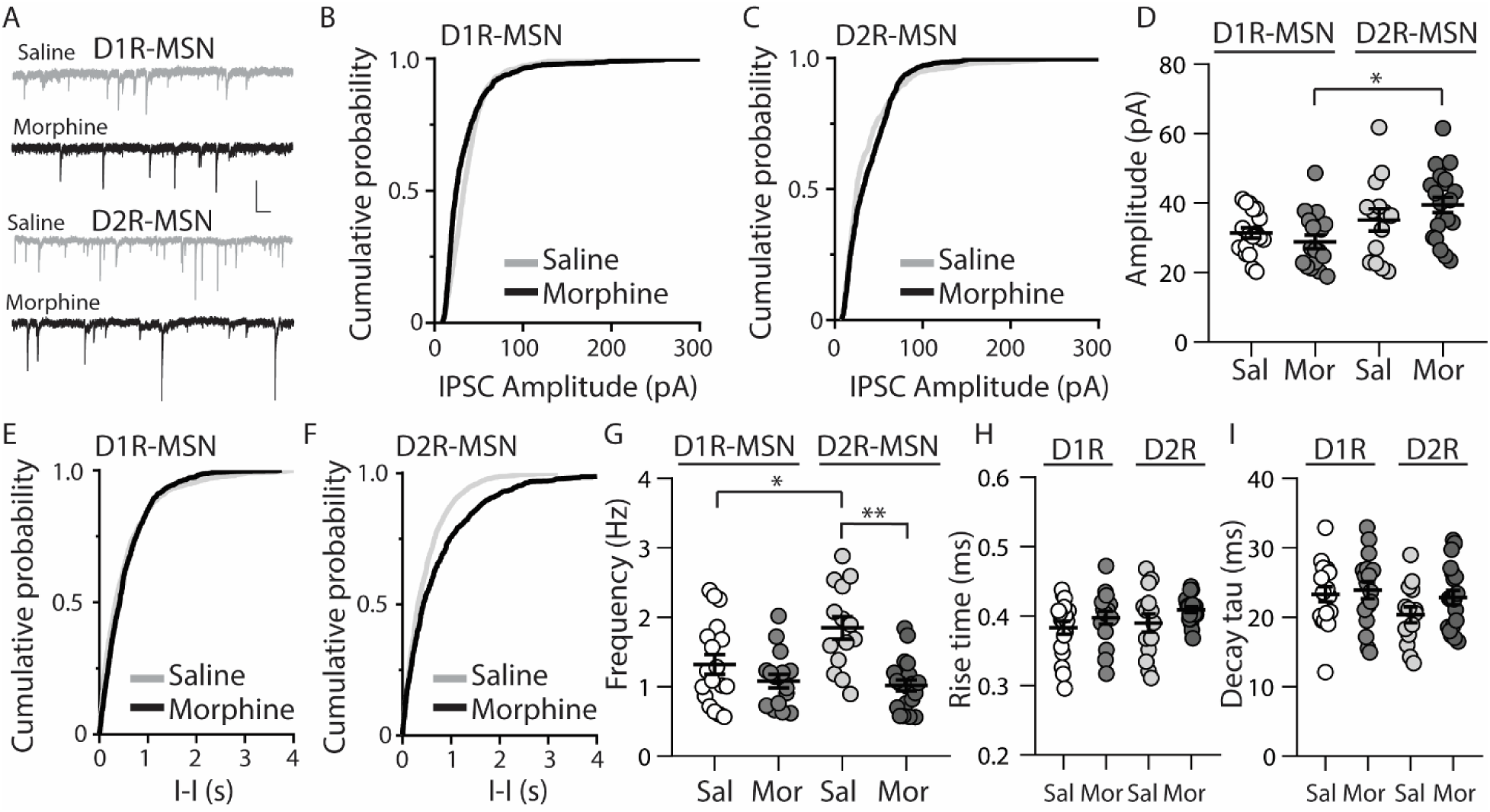
Repeated morphine administration reduces mIPSC frequency on D2R-MSNs on abstinence day 1. (A) Representative traces showing mIPSCs recorded from D1R- or D2R-MSNs from animals treated with saline or morphine (10 mg/kg, i.p.). Scale bar: 25 pA, 0.5 s. (B and C) Cumulative plot showing the distribution of mIPSC amplitudes recorded from D1R-MSNs (B) or D2R-MSNs (C) in animals treated with saline or morphine. (D) Summary graph showing the average mIPSC amplitude recorded D1R- or D2R-MSNs following saline or morphine treatment (F_(3,65)_=4.73, p=0.005; one way ANOVA with Bonferroni post-test). *p<0.05. (E and F) Cumulative plot of a representative neuron showing the distribution of mIPSC inter-event intervals (I-I) recorded from D1R-MSNs (E) or D2R-MSNs (F) in animals treated with saline or morphine. (G) Summary graph showing the average mIPSC frequency recorded from D1R- or D2R-MSNs following saline or morphine treatment (F_(3,65)_=8.94, p<0.0001; one-way ANOVA with Bonferroni post-test; *p<0.05, **p<0.01). (H) Summary graph showing the average rise time of mIPSC recorded from D1R- or D2R-MSNs following saline or morphine treatment. (I) Summary graph showing the average decay tau of mIPSC recorded from D1R- or D2R-MSNs following saline or morphine treatment. Circle=neuron.

### 3.2. Morphine increases the intrinsic membrane excitability of D2R-MSNs

MSNs in the nucleus accumbens shell display bistable membrane potential properties characterized by a hyperpolarized quiescent “down” state and a depolarized “up” state associated with neuronal discharge (O’Donnell et al., 1999). These states are controlled by combined excitatory synaptic discharge and intrinsic membrane excitability (Huang et al., 2011), which are posited to bring the membrane potential close to the MSN firing threshold, thus impacting the efficiency of information relay to downstream brain regions (O’Donnell and Grace, 1995;Ishikawa et al., 2009). Given our observed changes in synaptically-mediated excitatory and inhibitory transmission on D2R-MSNs (Figs. 1 and 3), our next experiment tested whether morphine impacts cell-type specific MSN intrinsic membrane excitability. To do this, using whole-cell electrophysiological recordings, we measured the number of action potentials in response to depolarizing currents, as this approach is often used to measure intrinsic membrane excitability (Desai et al., 1999;Nelson et al., 2003;Zhang and Linden, 2003;Heng et al., 2008;Ishikawa et al., 2009;Wang et al., 2018). We found that during morphine abstinence, there were no changes on D1R-MSN membrane excitability (Bonferroni post-test at each current injected, p>0.999) (Fig. 4A and B). However, the morphine-induced decreases in synaptic input onto D2R-MSNs (Figs. 1 and 3) were accompanied by an overall increase in the intrinsic membrane excitability at currents of ≥250 pA (Bonferroni post-test, 250 pA: p=0.008; 300 pA: p= 0.0003; 350-450 pA: p<0.0001) (Fig. 4A and C).

**Figure 4.**
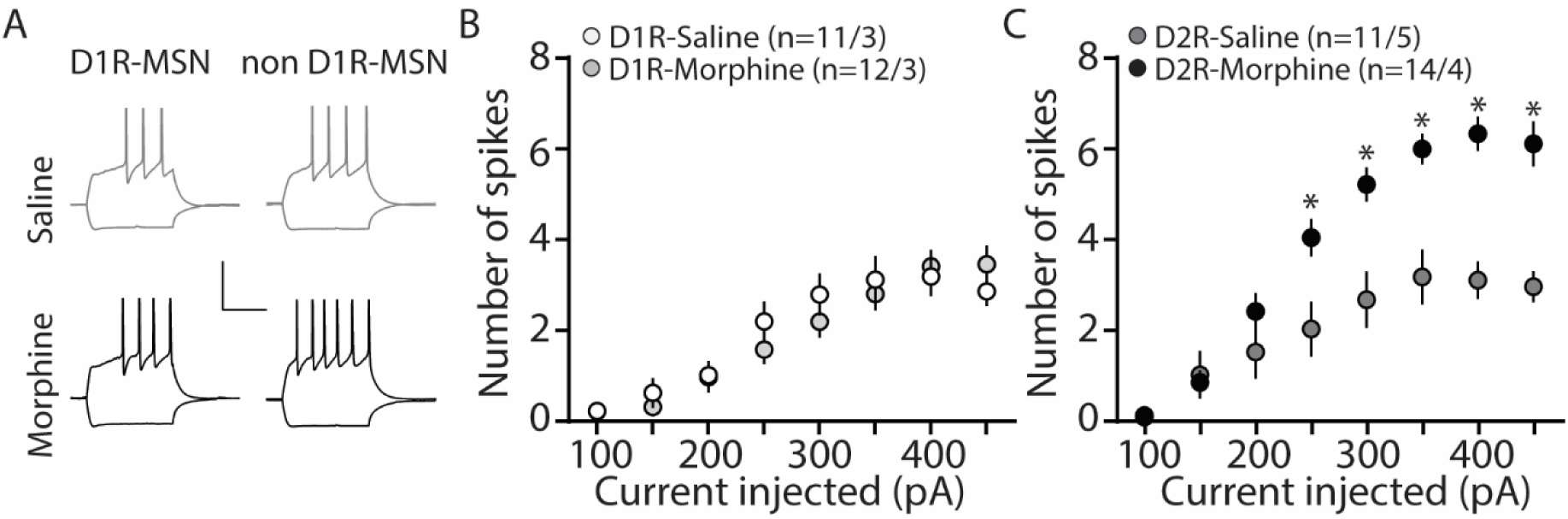
Repeated morphine administration increases membrane excitability on D2R-MSNs on abstinence day 1. (A) Representative traces, scale bar, 40 mV, 300 ms at 100 pA current injection. (B) Summary graph showing the average number spikes generated by injected current on D1R-MSNs following saline or morphine (10 mg/kg, i.p.) treatment (F_(7,238)_=1.05, p=0.395; two-way repeated measures ANOVA). (C) Summary graph showing the average number spikes generated by injected current on D2R-MSNs following saline or morphine treatment (F_(7,210)_=10.4, p=0<0.0001; two-way repeated measures ANOVA with Bonferroni post-test). *p<0.05. (n/n=cells/animals).

### 3.3. D2R-MSN synaptically driven functional output is unchanged following morphine treatment

Our present findings demonstrate that morphine exposure decreases mEPSC or mIPSC frequency and increases the intrinsic membrane excitability on D2R-MSNs. In an attempt to determine the integrated effect of these alterations on the overall functional output of D2R-MSNs, we measured the input-output efficacy by measuring synaptically-driven action potential firing (Hopf et al., 2003;Otaka et al., 2013). This was performed by counting the number of action potentials generated on D1R- or D2R-MSNs when varying currents (0-100 μA, 5 μA increments) were injected through a stimulating electrode during a 10 Hz stimulus. These measurements were performed in the absence of pharmacological blockers in the bath solution, thus cell-type specific MSN responses were influenced by mixed excitatory and inhibitory inputs (see Materials and Methods). Following morphine treatment, stimulating afferents in the nucleus accumbens elicited similar NBQX-sensitive action potential responses (Fig. 5A) in D1R- (Fig. 5B) or D2R-MSNs (Fig. 5C) compared to saline controls (D1R-MSN: F_(20,220)_=0.349, p=0.996; D2R-MSN: F_(20,260)_=1.05, p=0.409, two-way repeated measures ANOVA). Since neuronal excitability is not only influenced by the current intensity, but also by the temporal aspects of the current pulse, we constructed strength-duration curves whereby the electrically-evoked current was plotted over the electrically-evoked current duration (Fig. 6). By constructing this curve, we were able to observe increases or decreases in pre- and postsynaptic connections shown as steep or shallow decays in amplitude, respectively, as the pulse duration increases (Fröhlich, 2016). Once plotted, the rheobase, minimal electrically stimulated current required to elicit an action potential at an infinite pulse duration, and the chronaxie, an indication of neuronal excitability defined by the duration of the stimulus corresponding to twice the rheobase, were calculated. 24 h following morphine treatment, we found that the rheobase was not significantly different compared to control conditions (F_(3,19)_=0.048, p=0.986, one-way ANOVA) (Fig. 6B). Similarly, the chronaxie on D1R- or D2R-MSNs showed no significant change following morphine treatment (F_(3,19)_=0.8445, p=0.486, one-way ANOVA) (Fig. 6C). Overall, these results suggest that the morphine-induced decreases in synaptic transmission on D2R-MSNs are countered by increases in intrinsic membrane excitability, which together, enable D2R-MSNs to maintain basal levels of functional output in response to synaptic input.

**Figure 5.**
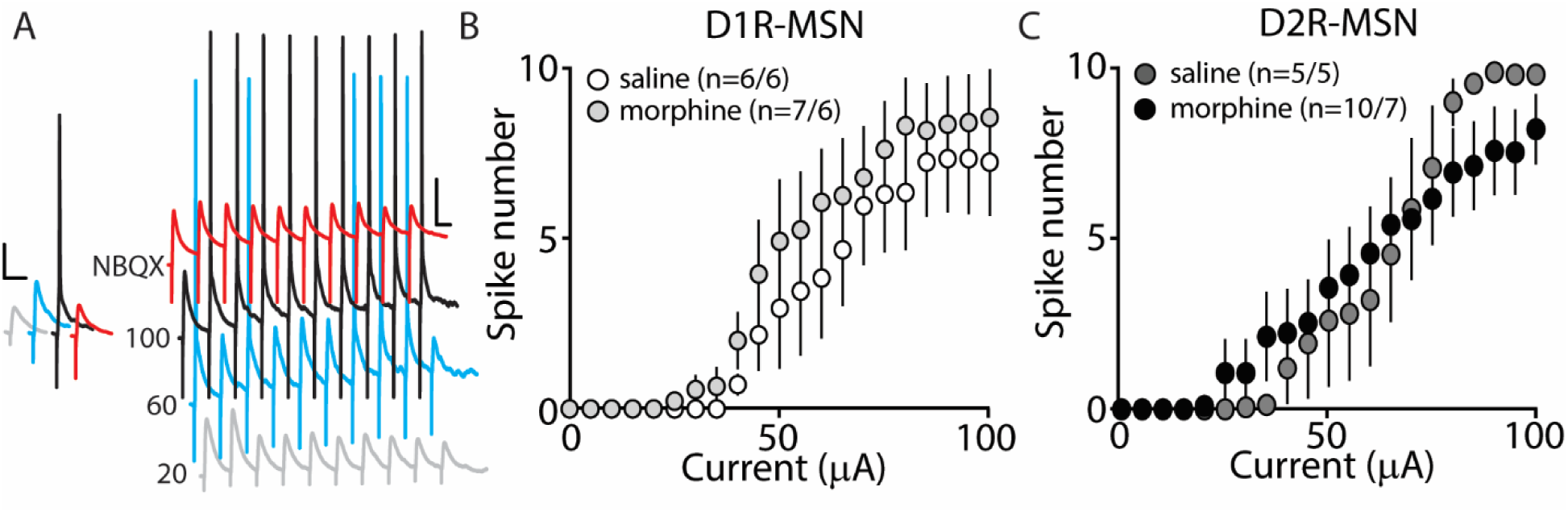
Synaptically-driven action potential firing on D1R- or D2R-MSNs is unaffected by repeated morphine (10 mg/kg, i.p.) treatment. (A) Representative traces showing depolarizations or action potentials of a recorded MSN evoked by electrical current (in μA) of 20 (light gray), 60 (blue), 100 (black), or 100 in the presence of NBQX (red), an AMPA receptor antagonist. Scale bars, 12.5 mV, 50 ms. (B and C) Summary graphs showing the average spike number at each current injected for D1R- or D2R-MSNs following saline or morphine treatment (cells/animals).

**Figure 6.**
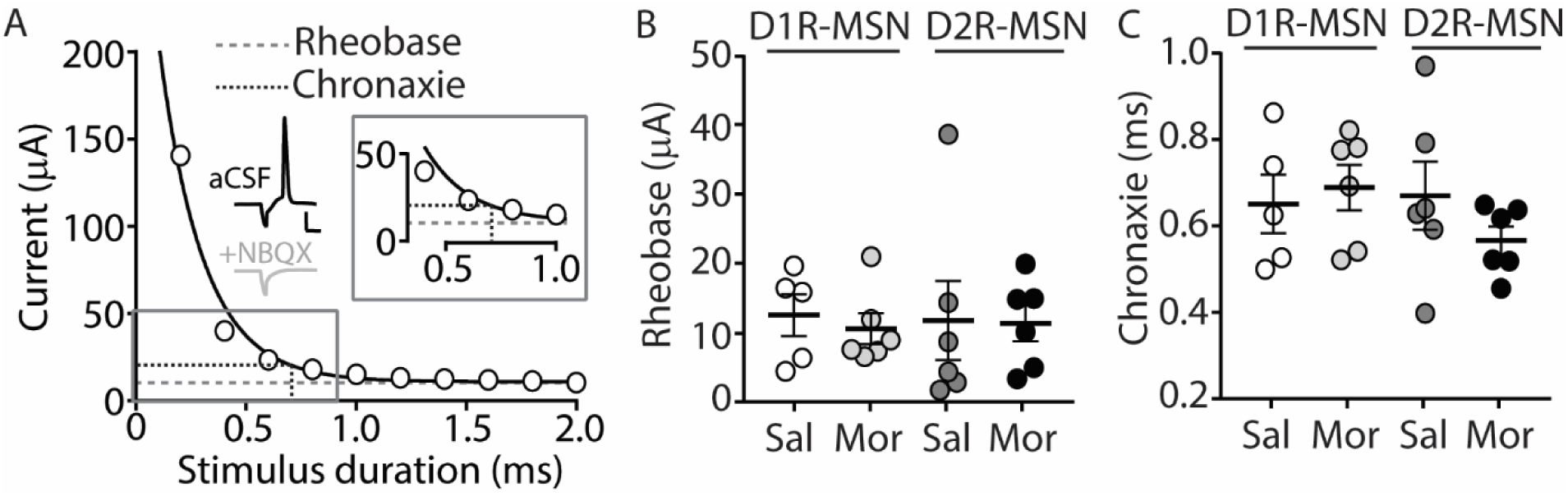
A strength-duration curve constructed from an MSN in the nucleus accumbens shell. Stimulus current was adjusted at each duration (from 0-2.0 ms with 0.2 ms increments) until an action potential was evoked using an electrical stimulus. The curve was fit with a two-phase exponential decay. The rheobase (gray dashed line) was calculated as the plateau of the curve and the chronaxie (black dashed line) was calculated as 2x the rheobase. (Inset) Representative traces illustrating the stimulus (downward deflection) followed by the action potential at 2 ms duration). In the presence of NBQX (2 μM), action potentials are not elicited (2 ms duration, 35 μA of current). Scale bars, 20 mV, 12.5 ms. (B) Summary graph showing the average rheobase for D1R- or D2R-MSNs following saline (Sal) or morphine (Mor) (10 mg/kg, i.p.) treatment. (C) Summary graph showing the average chronaxie for D1R- or D2R-MSNs following saline or morphine treatment.

## 4. Discussion

Our results show that repeated morphine administration preferentially alters action-potential independent synaptic transmission and intrinsic membrane excitability on D2R-MSNs, without affecting D1R-MSNs. Furthermore, our results show that the synaptically-driven action potential responses on D2R-MSNs, which are expected to integrate both synaptic and intrinsic cellular properties, remain unchanged following morphine exposure.

### 4.1. Morphine-induced changes in MSN intrinsic membrane excitability

A neuronal homeostatic response refers to a self-correcting property that is necessary in order to maintain stable function (Huang et al., 2011;Turrigiano, 2011). Here, our results show that 24 h post morphine treatment, the overall synaptic input on D2R-MSNs is reduced (Figs. 1 and 3), while the intrinsic membrane excitability is significantly increased (Fig. 4). Given that the mammalian central nervous system, including the nucleus accumbens, is capable of compensatory changes in intrinsic membrane excitability to overcome attenuated synaptic function (Burrone et al., 2002;Maffei and Turrigiano, 2008;Ishikawa et al., 2009), it is possible that a homeostatic synaptic-to-membrane crosstalk enables D2R-MSNs to maintain sensitivity to incoming signals, which is supported by our observed non-significant change in functional output following morphine exposure (Figs. 5 and 6).

This potential homeostatic response to morphine is in line with observations in the nucleus accumbens, during short-term abstinence from repeated cocaine administration, whereby NMDA receptor synaptic expression is increased (Huang et al., 2009), while in parallel, MSN intrinsic membrane excitability is decreased (Zhang et al., 1998;Dong et al., 2006;Ishikawa et al., 2009;Kourrich and Thomas, 2009;Mu et al., 2010;Wang et al., 2018). Although, both cocaine and morphine elicit homeostatic compensatory changes, the contrasting effects on MSN synaptic transmission and intrinsic membrane excitability is potentially due to the drug’s cell-type specific effects in the accumbens (Huang et al., 2009;Brown et al., 2011;Graziane et al., 2016).

Alternatively, homeostasis may not drive the opposing synaptic and intrinsic morphine-induced changes on D2R-MSNs as these changes may be two independent adaptations. Given that we observed morphine-induced reductions in both excitatory and inhibitory synaptic transmission on D2R-MSNs, the overall synaptic transmission may produce no net changes. This is supported by the sEPSC-to-sIPSC or the sIPCS-to-sEPSC inter-event intervals that show no change following morphine treatment (Figs. 2D and E). These results suggest that the increases in the intrinsic membrane excitability would result in an overall excitation gain on D2R-MSNs. Although, an excitation gain was not observed in our attempt to integrate both the morphine-induced synaptic and intrinsic properties (Figs. 5 and 6), it is possible that *in vivo* a more complicated scenario exists. Extensive evidence demonstrates that drugs of abuse influence the firing properties of neurons in the accumbens (Peoples et al., 1999;Carelli and Ijames, 2000;Ghitza et al., 2006;Calipari et al., 2016). Given that the bistable membrane potentials of MSNs (e.g., ~-80 mV down state versus ~-60 mV up state) (O’Donnell and Grace, 1995) are regulated by synaptic input and intrinsic factors (Plenz and Kitai, 1998;Wickens and Wilson, 1998;Huang et al., 2011), dysregulations in these factors may influence information flow from MSNs to downstream targets (O’Donnell et al., 1999), potentially influencing motivated behaviors.

Lastly, a previous study has shown that morphine exposure decreases the intrinsic membrane excitability on MSNs in the nucleus accumbens (NAc) with concomitant increases in the action potential amplitude, decreases in the action potential half-width, and decreases in the membrane resistance and tau (Heng et al., 2008). In contrast, following morphine exposure, we observed an increase in the intrinsic membrane excitability on D2R-MSNs in the NAc shell with no changes in action potential amplitude or half-width, increases in membrane resistance, and no changes in tau. These discrepancies may be a result of a number of differences between studies including the species (rat versus mice), the recording location (unspecific recordings in the NAc versus NAc shell), the exposure and recording paradigm (7 d morphine with recordings 3-4 d post treatment versus 5 d morphine with recordings 24 h post treatment), and/or the bath temperature during recordings (30-32°C versus 22-24°C). Regardless of this discrepancy, a key finding from our studies was the robust morphine-induced increase in intrinsic membrane excitability on D2R-MSNs, while the intrinsic membrane excitability on D1R-MSNs remained unaltered. These results demonstrate that morphine exposure produces cell-type specific alterations within the reward neurocircuit.

### 4.2. Excitatory-inhibitory balance

The spatiotemporal interaction between excitatory and inhibitory synaptic connections on a targeted neuron regulates neuronal activity (Wehr and Zador, 2003;Zerlaut and Destexhe, 2017;He and Cline, 2019), modulates neuronal oscillations (Buzsaki and Wang, 2012), and balances network dynamics (van Vreeswijk and Sompolinsky, 1996;Berke, 2009;Buzsaki and Watson, 2012;Deneve and Machens, 2016;Bonnefond et al., 2017). A problem arises when the E-I balance is disrupted causing a chronic deviation from the original set-point, which is associated with pathological states including autism, schizophrenia, epilepsy, and addiction-like behaviors (Rubenstein and Merzenich, 2003;Eichler and Meier, 2008;Fritschy, 2008;Yizhar et al., 2011;Tejeda et al., 2017;Yu et al., 2017). Here, we investigated whether morphine abstinence alters the E/I ratio on D1R- or D2R-MSNs in the accumbens shell by comparing the evoked excitatory to inhibitory current amplitudes as well as the temporal integration of spontaneous excitatory and inhibitory events. We show that despite the morphine-induced changes in mEPSC and mIPSC frequency on D2R-MSNs, the E/I evoked current amplitude ratio and the temporal relationship between spontaneous excitatory and inhibitory events were unchanged following morphine administration (Fig. 2), potentially due to homeostatic mechanisms that tightly maintain neuronal E-I balance on D2R-MSNs (Turrigiano and Nelson, 2000;2004;Turrigiano, 2011).

We have not examined the mechanisms triggering the potential homeostatic mechanisms that maintain the E-I balance. However, previously, it has been shown that morphine-induced decreases in glutamatergic transmission on D2R-MSNs are prevented by administration of the GluA_2_3Y peptide, which prevents morphine-induced AMPAR removal from excitatory synapses (Ahmadian et al., 2004;Brebner et al., 2005;Wang, 2008;Graziane et al., 2016;Madayag et al., 2019). Future studies can investigate the synaptic cascade that potentially leads to the maintenance of the E-I balance on D2R-MSNs following morphine exposure, by administering GluA_2_3Y peptide and measuring effects on D2R-MSN mIPSC frequency and intrinsic membrane excitability. Such work may reveal a homeostatic mechanism triggered by morphine-induced decreases in glutamatergic synaptic transmission that may also regulate intrinsic membrane excitability.

Lastly, we observed a non-significant change in functional output on D2R-MSNs following morphine exposure (Figs. 5 and 6) despite the increases in intrinsic membrane excitability (Fig. 4). This result is potentially explained by our synaptic assessments showing an overall decrease in mEPSC frequency, mIPSC frequency, with no change on the mEPSC or mIPSC amplitude, E/I ratio, or on the temporal relationship between the E-I balance (Fig. 2). These results suggest that, following morphine exposure, the overall somatic summation of excitatory and inhibitory currents on D2R-MSNs is potentially weakened. This is likely caused by weakened postsynaptic excitatory glutamatergic synaptic connections (i.e., decreases in mEPSC frequency (Fig. 1) and increases in the expression of silent synapses (Graziane et al., 2016)) as well as the alterations in presynaptic factors that result in decreased inhibitory synaptic transmission (i.e., decreases in mIPSC frequency (Fig. 3)). These morphine-induced decreases in synaptic transmission on D2R-MSNs along with the morphine-induced increases in intrinsic membrane excitability, together, likely enable D2R-MSNs to maintain basal levels of functional output in response to synaptic input.

### 4.3. Receptor kinetics

AMPA/kainate receptor kinetics comprise a rapidly rising conductance that decays as the agonist-receptor complex deactivates (Traynelis et al., 2010). This process is regulated by a number of factors including receptor subunit composition (Sommer et al., 1990;Partin et al., 1996;Quirk et al., 2004) and auxiliary regulatory proteins (Milstein et al., 2007;Milstein and Nicoll, 2008). Here, we show that 24 h post morphine treatment, the AMPA/kainate receptor kinetics (rise time and decay tau) on D1R- or D2R-MSNs are unchanged (Fig. 1H and I). This result cannot exclude potential alterations in morphine-induced auxiliary protein expression or receptor subunit composition. It has been shown that mRNAs for AMPA receptor subunits GluA1, 3, and 4 are significantly decreased in morphine self-administering rats (Hemby, 2004). However, on average, any morphine-induced changes that may occur, are unable to elicit changes in overall kinetic properties of AMPA/kainate receptors responding to action potential independent glutamate release. This suggests that if morphine-induces any potential changes in EPSC temporal summation at the soma, these alterations are likely not mediated by changes in AMPA/kainate receptor kinetics. Similarly, we observed no changes in inhibitory ionotropic neurotransmitter receptor kinetics on D1R- or D2R-MSNs following morphine treatment (Fig. 3H and I), again suggesting that, overall, if morphine was able to induce changes in receptor phosphorylation, density, or scaffolding proteins, factors regulating inhibitory ionotropic receptor kinetics (Verdoorn et al., 1990;Takahashi et al., 1992;Tia et al., 1996;Jones and Westbrook, 1997;Chen et al., 2000), they are unable to influence the overall kinetic properties of inhibitory ionotropic receptors responding to action potential independent neurotransmitter release.

### 4.4. Sex comparisons

We found that, within all measurements where male and female mice were used (e.g., mEPSC recordings, mIPSC recordings, E/I ratios, intrinsic membrane excitability, synaptically-driven action potentials, and rheobase/chronaxie measurements), there were no statistically significant sex differences within D1R- or D2R-MSNs following non-contingent, repeated saline or morphine treatment (**Table III**). Because of this, animals were pooled. However, we understand that our statistical assessment is likely underpowered and therefore, future experiments are required to directly test sex differences. Additionally, it is possible that bimodal distributions in our data set are influenced by sex effects. For example, Fig 1G shows a bimodal distribution in the D1R-MSN cell population following morphine treatment. However, upon further analysis, these two populations consist of neurons from both males and females suggesting that at the 24 h abstinence time point following repeated morphine administration, mEPSC frequency on D1R-MSNs is unaltered. Despite this, it is still worthwhile to perform a thorough assessment of potential sex effects as it has been shown that, under basal conditions, D2R-MSN mEPSC frequency is significantly reduced in female versus male prepubertal (2-3 week old) mice, in the accumbens core (Cao et al., 2018). Determining whether these sex differences are observed into adulthood following morphine exposure would be an interesting future direction.

**Table III.** Sex comparisons within electrophysiological assessments investigating morphine-induced changes in synaptic and intrinsic properties of D1R- or D2R-MSNs.

| Experiment | Saline | D1R | P Value | Saline | D2R | P Value | Morphine D1R |  | P Value | Morphine D2R |  | P Value |
| --- | --- | --- | --- | --- | --- | --- | --- | --- | --- | --- | --- | --- |
|  | Male | Female |  | Male | Female |  | Male | Female |  | Male | Female |  |
| **mEPSC Frequency (Fig. 1)** | 6.67±0.51 (6) | 6.82±0.68 (6) | 0.86 | 6.32±1.43 (5) | 5.55±1.40 (7) | 0.71 | 5.17±0.55 (4) | 4.61±0.61 (7) | 0.58 | 3.33±0.44 (8) | 2.30±0.28 (3) | 0.21 |
| **mEPSC Amplitude (Fig. 1)** | 15.12±1.34 (6) | 14.34±1.14 (6) | 0.67 | 12.48±1.10 (5) | 13.55±1.07 (7) | 0.51 | 14.58±1.29 (4) | 14.15±0.83 (7) | 0.78 | 12.28±0.56 (8) | 11.50±0.51 (3) | 0.46 |
| **E/I Ratio (Fig. 2)** | 0.22±0.06 (2) | 0.37±0.10 (9) | 0.50 | 0.28±0.05 (2) | 0.46±0.14 (7) | 0.52 | 0.25±0.10 (3) | 0.38±0.09 (8) | 0.47 | 0.09±0.03 (2) | 0.32±0.06 (8) | 0.12 |
| **mIPSC Frequency (Fig. 3)** | 1.10±0.17 (5) | 1.41±0.18 (13) | 0.35 | 1.55±0.21 (4) | 1.97±0.20 (10) | 0.26 | 1.21±0.15 (5) | 1.03±0.12 (12) | 0.43 | 0.91±0.12 (4) | 1.05±0.10 (16) | 0.50 |
| **mIPSC Amplitude (Fig. 3)** | 31.44±2.46 (5) | 31.39±1.94 (13) | 0.99 | 34.20±3.91 (4) | 35.57±4.30 (10) | 0.85 | 30.62±2.09 (5) | 28.09±2.63 (12) | 0.57 | 33.27±4.57 (4) | 41.07±2.47 (16) | 0.17 |
| **IME (450pA) (Fig. 4)** | 2.43±0.20 (7) | 2.75±0.48 (4) | 0.49 | 4.29±0.97 (7) | 2.50±0.29 (4) | 0.21 | 2.86±0.40 (7) | 4.20±1.36 (5) | 0.30 | 5.67±0.88 (6) | 6.38±0.65 (8) | 0.52 |

Number of cells (n). Student's t-test was used for statistical measures. Abbrev.: Miniature excitatory postsynaptic current, mEPSC; excitatory to inhibitory ratio, E/I ratio; miniature inhibitory postsynaptic current, mIPSC; intrinsic membrane excitability, IME.

### 4.5. Cell type comparisons

The nucleus accumbens is a complicated network consisting of D1R and D2R-MSNs that project to similar brain regions (Smith et al., 2013). Additionally, it has been shown that lateral inhibition between MSNs exists and this lateral inhibition is critically involved in addiction-like behaviors (Dobbs et al., 2016). Therefore, imbalances in D1R- and D2R-MSN activity, in downstream targets, or within the accumbens microcircuit, are potentially responsible for behavioral phenotypes (e.g., locomotor activity or conditioned place preference) observed following repeated morphine treatment (Zarrindast et al., 2002;Bohn et al., 2003;Tzschentke, 2007). Using our statistical approach, we found a significant difference in mIPSC amplitude between D1R-MSN morphine and D2R-MSNs morphine (F_(3,65)_=4.73, p=0.005; one way ANOVA with Bonferroni post-test revealing significant differences between D1R-MSN morphine and D2R-MSN morphine, p=0.0048) (Fig. 3D). This result provides a potentially interesting opportunity to determine whether significant differences in electrophysiological readouts between neuronal types is sufficient to contribute to drug-induced behavioral phenotypes. For example, increasing D1R-MSN mIPSC amplitude or decreasing D2R-MSN mIPSC amplitude in morphine-treated animals may block morphine-induced behavioral phenotypes as cell-type interactions may drive morphine-induced behaviors. This idea may also be applied to our observed significant difference in mIPSC frequency between D1R- and D2R-MSNs in saline-treated animals (p=0.024, Bonferroni post-test), which was non-significant following morphine treatment (Fig. 3G). Based on these cell-type specific comparisons in electrophysiological data, it will be interesting to test whether cell-type specific interactions significantly contribute to addiction-like behaviors.

## 5. Conclusions

In conclusion, this study demonstrates new information on how morphine exposure alters both extrinsic and intrinsic neuronal properties of MSNs in the nucleus accumbens shell. The alterations observed on D2R-MSNs appear to be opposing in nature, resulting in a maintenance of basal levels of functional output. It is known that preventing morphine-induced decreases in glutamatergic transmission on D2R-MSNs blocks the prolonged maintenance (21 d post conditioning) of morphine-induced CPP (Graziane et al., 2016). Therefore, it is plausible that morphine-induced alterations on synaptic and intrinsic excitability of D2R-MSNs may not alter D2R-MSN output during short-term abstinence, but may instead result in an allostatic set point of excitability that results in long-term behavioral consequences (Koob and Le Moal, 2001). Although, future studies are required to directly test whether the observed maintenance of D2R-MSN output drives the prolonged expression of opioid-seeking behaviors.

## Acknowledgements

We thank Dr. Diane McCloskey for edits and comments, and the Silberman lab for their comments on the project. The study was supported by the NARSAD Young Investigator Award (27364-NG), the Pennsylvania State Junior Faculty Scholar Award (NG), the Pennsylvania Department of Health using Tobacco CURE Funds (NG), and the Pennsylvania State Research Allocation Project Grant (NG). Morphine was provided by the Drug Supply Program of NIDA NIH.

## Author Contributions Statement

D.S.M. and N.M.G. designed the experiments and analyses, conducted the experiments and data analyses, and wrote the manuscript. B.J. designed a program for the analysis performed in Fig. 2.

